# Matching Excellence: ONT’s Rise to Parity with PacBio in Genome Reconstruction of Non-Model Bacterium with High GC Content

**DOI:** 10.1101/2024.02.26.582104

**Authors:** Axel Soto-Serrano, Wenwen Li, Farhad M. Panah, Yan Hui, Pablo Atienza, Alexey Fomenkov, Richard J. Roberts, Paulina Deptula, Lukasz Krych

## Abstract

Reconstruction of complete bacterial genomes is a vital aspect of microbial research, as it provides complex information about genetic content, gene ontology, and regulation. It has become a domain of 3rd generation, long-read sequencing platforms, as short-read technologies can deliver mainly fragmented genomes. PacBio platform can provide high-quality complete genomes, yet remains one of the most expensive sequencing strategies. Oxford Nanopore Technology (ONT) offers the advantage of producing the longest reads, being at the same time the most cost-effective option in terms of platform costs, as well as library preparation, and sequencing. However, ONTs error rate, although significantly reduced lately, still holds a certain level of distrust in the scientific community.

In recent years, hybrid assembly of Nanopore and Illumina data has been used to solve ONTs issue with error rate and has yielded the best results in terms of genome completeness, quality, and price. However, the latest advancements in Nanopore technology, including new flow cells (R10.4.1), new library preparation chemistry (V14) and duplex-mode, updated basecallers (Dorado v0.4.1), and the realization that sequencing in dark mode results in significantly increased throughput, have had a significant impact on the quality of generated data and, thus, the recovery of complete genomes by ONT sequencing alone.

In this study, we compared the data generated by ONT using three sequencing strategies (Native barcoding, RAPID barcoding, and custom-developed: BARSEQ) against PacBio and Illumina (NextSeq) as well as Illumina-ONT hybrid data. For this purpose, we employed three strains of the actinobacteria *Propionibacterium freudenreichii*, whose genomes have been proven difficult to reconstruct due to high GC content, regions of repeated sequences and massive genome rearrangements.

Our data indicate that DNA libraries prepared with the native barcoding kit, sequenced with V14 chemistry on R10.4.1 flow cell, and assembled with Flye resulted in the reconstruction of complete genomes of overall quality highly similar to that of genomes reconstructed with PacBio. The highest level of quality can be achieved by hybrid assembly of data from the Native barcoding kit complemented with data from custom-developed BARSEQ, both sequenced on R10.4.1 flow cell. In conclusion, our results demonstrate that ONT can be used as a cost-effective sequencing strategy, without the need for complementing with other sequencing technologies, for the reconstruction of complete genomes of the highest quality.

## Introduction

The reconstruction of complete, high-quality bacterial genomes is crucial for understanding bacterial diversity, evolution, and ecological roles. Studying full bacterial genomes provides crucial insights into the basis of pathogenicity, antibiotic resistance, and other virulence factors (Ghazi et al., 2022; Zhao et al., 2023). For the past two decades, next-generation sequencing (NGS) platforms have dominated and revolutionized microbial genomics research, allowing accurate and cost-effective sequencing. However, their main limitation, short-read sequences, often hinders the recovery and assembly of complete, circular genomes. For example, most commonly used Illumina technology allows high accuracy reads (>99%), but only of short fragments of 500bp or fewer (Wick et al., 2017), compromising adequate full genome assembly when repeat regions are larger than such fragments (Achaz et al., 2002; Siguier et al., 2006).

The advent of third generation sequencing platforms capable of producing long reads, such as Pacific Biosciences (PacBio) and Oxford Nanopore Technology (ONT), enabled reconstruction of complete genomes, though initially both technologies suffered from significantly high error rates (∼ 10-15%;Eid et al., 2009; Ip et al., 2015) on a single molecule level, severely affecting the quality of recovered genomes. PacBio was the first 3^rd^ Generation Sequencing that has largely improved the quality, reaching today 99.999% consensus accuracies according to the manufacturer, and although recovery of good quality complete genomes became possible on a single platform, the data generation with this technology remains very costly.

ONT struggled longer with the error on a single molecule level. In this context, hybrid assembly utilizing tools such as Unicycler (Wick et al., 2017) has been a vastly used solution to obtain complete and accurate sequences. This approach involves polishing long reads obtained with ONT with inexpensive and accurate Illumina short read data. Despite enduring challenges, great efforts have been made by ONT to reduce error rates, and today the company reports accuracies close to 99% (Marx, 2023). Only in 2023, ONT has introduced important upgrades in chemistry (V14), flow cells (R10.4.1), duplex mode, and new basecalling algorithms (Dorado v0.5.1).

Nanopore offers the cheapest sequencing platform on the market and a very broad portfolio of flow cells and library preparation strategies. Furthermore, it allows DNA sequencing and library preparation without PCR, enabling to study base methylations, and it generates by far the longest reads reaching megabases (Wang et al., 2021). Moreover, recent (August 2023) news about the light sensitivity of V14 chemistry led to instant adjustment in all protocols to prepare, load, and sequence ONT libraries in darkness. This was speculated to improve the throughput of long read sequencing up to (141%) and amplicon sequencing even up to (469%) (Vilella, 2023), though official peer-review data on that and whether it could also affect the quality are not available at this time.

Despite many advantages of nanopore technology, it remains unclear as to what extent different library preparation strategies, newest chemistry as well as flow cell have in fact reduced the error rate. In this paper, we compared sequencing data previously generated on Illumina NextSeq platform, ONT R9.4.1, and PacBio as gold standard, to data generated by newest ONT flow cells R10.4.1, utilizing different library preparation (Native barcoding, RAPID barcoding, and custom developed: BARSEQ) and assembly strategies for three strains of the beneficial actinobacteria *Propionibacterium freudenreichii.* The genomes of these bacteria have been proven difficult to reconstruct due to high GC content, regions of repeated sequences and massive genome rearrangements (Deptula et al., 2017; Ojala et al., 2017). To assess resulting assemblies, we used CheckM-based genome completeness, whole genome variant calling, gene annotation comparisons, and methylation motif recovery.

## Materials and Methods

### 1. Bacteria growth and genomic DNA extraction

*Propionibacterium freudenreichii* TL110, TL19, and TL29 were streaked from glycerol (15%) stocks and first grown anaerobically (Anaerogen) on Yeast Extract Lactate (YEL) agar for 4 days and then in YEL broth for 48h, both at 30°C. The grown bacterial cells were subsequently centrifuged at 5000 G for 10 min using a 5920 R centrifuge (Eppendorf, Hamburg, Germany) washed in 1 mL 0.9% NaCl and subsequently centrifuged again at 12000 rpm for 3 min. The resulting cell pellets were used for extraction of genomic DNA with a Bead-Beat Micro AX Gravity kit following the manufacturer’s instructions (cat # 106-100-M1, A&A Biotechnology, Gdynia, Poland) for ONT sequencing on R10.4.1 flowcells, and with MagAttract HMW DNA kit (Qiagen, Germantown, MD, USA) for Illumina, PacBio and ONT sequencing with R9.4.1 flow-cells. Genomic DNA was quantified using a Qubit 4 fluorometer (Thermofisher, Waltham, USA). A summary of the experimental workflow including these, and the following steps in this study, is presented in Figure 1.

**Figure 1.**
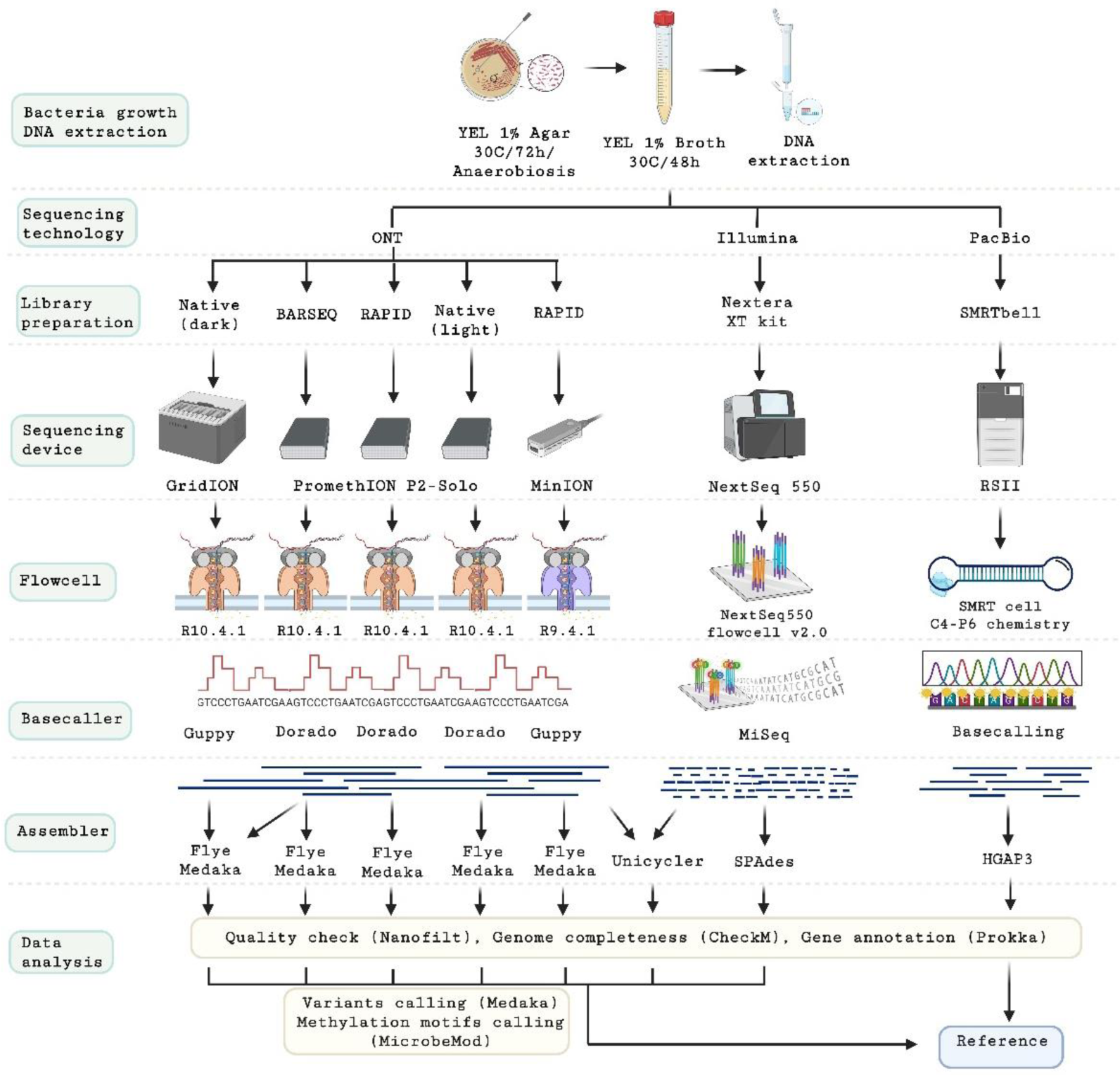
Workflow of the experiments. Figure created with BioRender.

### 2. DNA library preparation and sequencing

#### 2.1 RAPID barcoding for ONT

DNA concentration was normalized to 20 ng/μl (200ng total) and subjected to library preparation using Rapid Barcoding Sequencing protocol following the manufacturer’s instructions (SQK-RBK114.24, version: RBK_9176_v114_revG_27Nov2022, ONT, Oxford, UK). The sequencing was performed on PromethION 2 Solo sequencing platform connected to custom build computing station (MinKNOW 21.05.20 and basecalled with Dorado v0.5.0, ONT, Oxford, UK).

For R9.4.1 RAPID sequencing, DNA concentration was normalized to 53.3 ng/μl (400 ng total) and subjected to library preparation using Native Barcoding Sequencing protocol (SQK-RBK004, version: RBK_9054_v2_revK_14Aug2019, ONT, Oxford, UK), following the manufacturer’s instructions. The sequencing was performed on MinION platform (ONT, Oxford, UK). Data acquisition was performed with MinKNOW 23.07.15.

#### 2.2 Native barcoding for ONT

For R10.4.1 Native sequencing, DNA concentration was normalized to 33.33 ng/μl (400 ng total) and subjected to library preparation using Native Barcoding Sequencing protocol (SQK-NBD114.24, version: NBE_9169_V144_revN_15Sep2022 (dark)) (Oxford Nanopore Technologies, Oxford, UK), following the manufacturer’s instructions. The sequencing was performed on GridION and PromethION 2 Solo platforms (ONT, Oxford, UK), utilizing R10.4.1 flow cells. Data acquisition was performed with MinKNOW 21.05.20. Duplex reads were selected using Duplex-Tools (https://pypi.org/project/duplex-tools/v0.3.3); both pairs and split pairs were further basecalled using Dorado v0.5.1.

#### 2.3 Barcode-Amplified Random Sequencing (BARSEQ)

DNA concentration was normalized to 50 ng/μl and sheared to 6 Kb fragments using Megaruptor® 2 (Diagenode, Seraing, Belgium). Shearing quality was estimated with TapeStation 4200 (Agilent Technologies, CA, USA). Sheared DNA was subjected to the ligation with palindromic adapters and PCR amplification using primers with custom barcodes. The sequence of the adapter and barcodes is given in the Supplementary Table 1. Sheared DNA (25 μl of concentration 0,5 ng/μl) was mixed with 1.75 μl of Ultra II End-prep Reaction Buffer and 1.5 μl of Ultra II End-prep Enzyme Mix (cat # E7546, New England Biolabs MA, USA) and incubated at 20°C for 5 min. and 65°C for 5 min. After the End-prep reaction the DNA was purified with 30 μl of AMPure XP beads according to manufactures manual (Beckman Coulter Genomic, CA, USA) and resuspended in 11 μl of nuclease-free water.

The custom double stranded adapter containing primer-binding site was ligated by mixing 10 μl of NEB TA/Blunt Ligase Master Mix (cat #M0367, New England Biolabs MA, USA), 2 μl of duplex adapter (1 μM) and 10 μl of End-prepped DNA, and incubated at room temperature for 10 min. The sequence of the duplex adapter is as follows, first strand: 5’(5phos)-CCGAGCCCACGAGAC-3’, second strand: 5’-GTCTCGTGGGCTCGGT-3’. After the ligation, the reaction was purified with AMPure XP beads according to manufactures manual (Beckman Coulter Genomic, CA, USA) and resuspended in 12 μl of nuclease-free water.

Finally, the DNA containing the ligated primer binding site was subjected to the PCR reaction. The PCR mix composed of 12 µl of PCRBIO Ultra Mix (PCR Biosystems Ltd, London, United Kingdom), 2 µL of barcode primers (5 µM, see Supplementary Table 1 for details), 11 µL of purified DNA with adapters. The PCR thermal conditions were as follows: denaturation at 95 °C for 3 mins; 18 cycles of 95 °C for 30 s, 55 °C for 30 s, 69 °C for 5 min., and final elongation at 72 °C for 4 min. The final PCR2 products were pooled and purified using AMPure XP beads (Beckman Coulter Genomic, CA, USA). The pooled, barcoded DNA was purified with AMPure XP beads (Beckman Coulter Genomic, CA, USA) according to manufactures manual in 1:1 proportions and resuspended in 20 μl of nuclease-free water. Lastly, purified amplicons were subjected to 1D genomic DNA library preparation by ligation protocol (SQK-LSK114, ONT, Oxford, UK) to complete library preparation for GridION (ONT, Oxford, UK) sequencing. ∼0.2 μg of amplicons were used for the initial step of end-prep. Approximately 40 ng of prepared amplicon library was loaded on a R10.4.1 flow cell. Data acquisition was performed with MinKNOW 21.05.20 and basecalled with Dorado v0.5.0. It is important to note that a similar protocol (ligation sequencing v14 - low input by PCR: SQK-LSK114) is now commercially available from ONT, although in our protocol barcoding takes place during PCR and final ligation is performed on pooled library (Figure 2).

**Figure 2.**
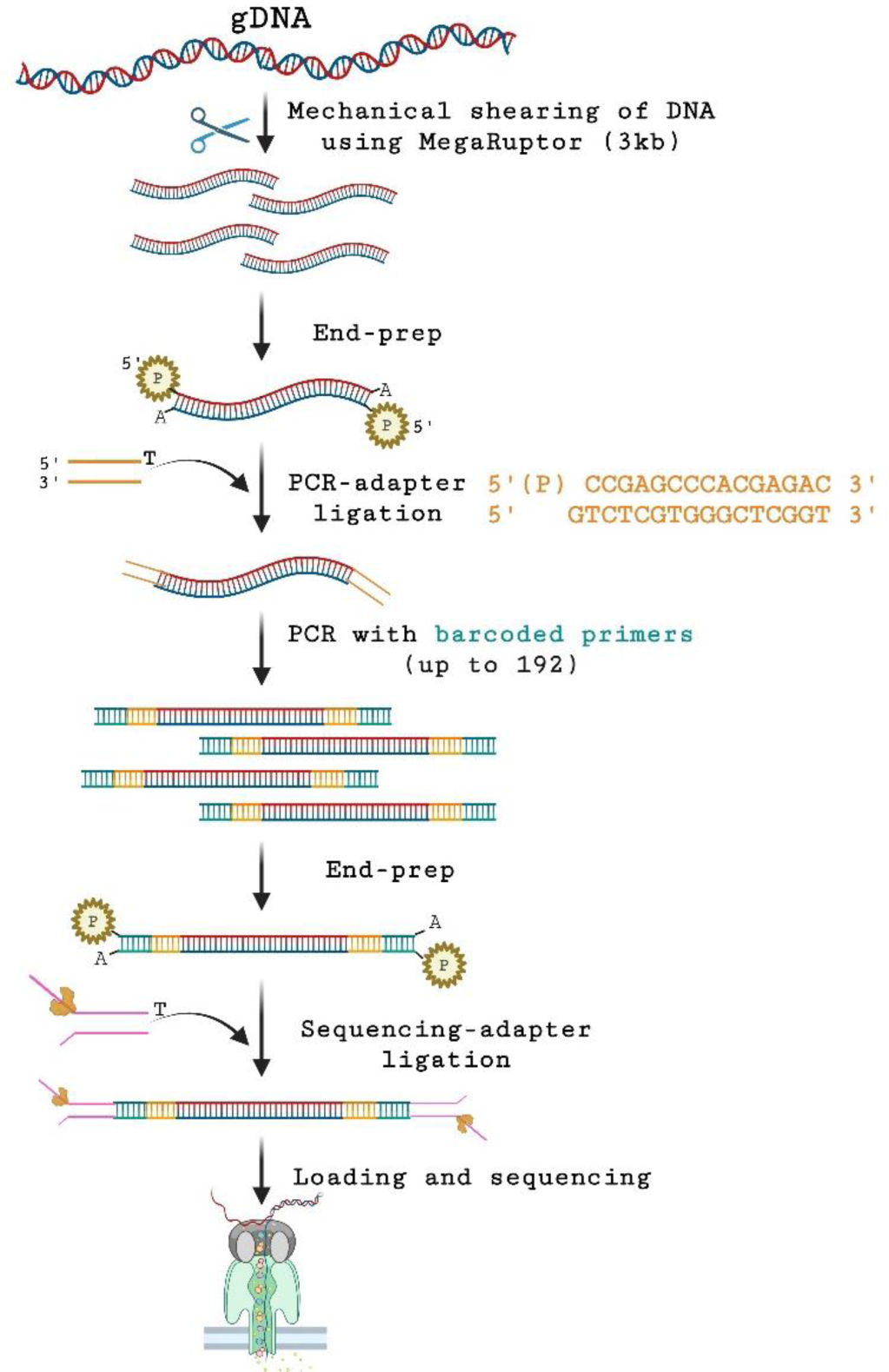
Overview of the BARSEQ protocol. Figure created with BioRender.

#### 2.4 Library preparation and sequencing for PacBio

DNA samples were sheared to an average size of ∼ 15 kb using the G-tube protocol (Covaris, MA). DNA libraries were prepared using a SMRTbell express template prep kit 2.0 (100-938-900, Pacific Bioscience, CA) and ligated with hairpin adapter. Incompletely formed SMRTbell templates were removed by digestion with a combination of exonuclease III and exonuclease VII (NEB, MA). The qualification and quantification of the SMRTbell libraries were made on a Qubit fluorimeter (Invitrogen, OR) and a 2100 Bioanalyzer (Agilent Technologies, CA). Additional separation of SMRTbell libraries on gel-based Blue Pippin instrument (Sage Science, Beverly, MA) was required due to inhibition of sequencing resulting from suspected contamination with high molecular weight carbohydrate polymers. Finally, SMRT sequencing was performed using a PacBio RSII (Pacific Biosciences, CA) sequencer with C4-P6 chemistry (7-10 SMRT cells with 300-minute collection time) based on protocol for 20 kb SMRTbell library inserts.

#### 2.5 Nextera tagmentation for Illumina

DNA concentration was normalized to 1 ng/μl and subjected to library preparation using Nextera XT DNA Library Preparation Kit (Illumina, CA, USA) according to the manufacturer’s instructions and sequenced with NextSeq550 using Mid Output Kit v2.5 (300 cycles, #20024905, Illumina, CA, USA), using a NextSeq550 v2.0 flow cell.

### 3. Data analysis

#### 3.1. Raw data quality

Quality scores of Illumina and ONT reads were assessed utilizing FastQC (v0.12.1) and a custom function (https://github.com/farhadm1990/ONT_helper), respectively. ONT reads were further filtered using Nanofilt (v2.6.0). The R10.4.1 and R9.4.1 reads were filtered by a quality score 16 and length 1000, and a quality of 7 and length 1000, respectively. Duplex reads for R10.4.1 Native were filtered by a quality score of 25 and a length of 1000.

#### 3.2. Read assembly

ONT reads were assembled using Flye (Kolmogorov et al., 2019), which has been encountered to be the best *de novo* assembler for long reads (Boostrom et al., 2022). In order to avoid any bias due to difference in reading depth for different ONT chemistries, all reads were normalized to 80X coverage for a genome size of 2.7 mbp, except for strain TL19 in R9.4.1 RAPID and R10.4.1 Rapid with 40 and 37X coverage, respectively. Flye v2.9.1 was used with default settings for all R9.4.1 and R10.4.1 samples. All the assembled contigs were subsequently polished by Medaka v1.11.0 (https://github.com/nanoporetech/medaka) and the resulting polished fasta file was used for downstream analysis.

Illumina Nextera 500 reads were assembled using SPAdes v3.15.4 (Bankevich et al., 2012) with the --isolate flag for higher sensitivity.

Hybrid assembly of ONT and Illumina reads was performed with Unicycler v0.5.0 (Wick et al., 2017) with conservative mode. It is noteworthy that the default assembler of Unicycler for ONT is miniasm+Racon (Li, 2016; Vaser et al., 2017), thus the ONT assemblies may differ from those obtained using Flye.

Finally, PacBio reads were *de novo* assembled using the RS_HGAP_Assembly3 version 2.3.0 program (Chin et al., 2013)with default quality and read length parameters.

All genomes were fixed for their replication start point based on *dnaA* gene using Circlator v1.5.5 and genomes with successfully fixed start were considered complete and circular.

#### 3.3. Gene annotation and genome completeness

All the assembled genomes were annotated using Prokka v1.14.6 (Seemann, 2014), considering hits with at least 80% similarity against its known protein database. Additionally, genome completeness within the known gene sets of the *Propionibacterium* genus was assessed using CheckM v1.2.2 (Parks et al., 2015).

The prevalence of unique genes across different sequencing strategies for the three strains was determined using a contingency matrix derived from the annotated genes. The contingency analysis excluded all shared genes, and the columns and rows were classified using the Ward.D2 distance method. Contingency heatmaps and dendrograms were generated using an R script (https://github.com/farhadm1990/ONT_helper).

#### 3.4. Variant calling

Variant calling, i.e., insertions and deletions (indels), as well as substitutions, including transitions and transversions was done against PacBio assemblies as reference. Variants calling was performed using medaka_haploid_variant command in Medaka. Only variants with a minimum Q score of 20 were selected for further analysis.

#### 3.5 Methylation calling

Methylation calling for ONT data was performed using MicrobeMod (Crits-Christoph et al., BioRxiv.) from Native R10.4.1 data. Basecalling was performed by dorado v0.5.1 utilizing the sup mode i.e., dna_r10.4.1_e8.2_400bps_sup@v4.3.0 with the --modified-bases-models flag using the models dna_r10.4.1_e8.2_400bps_sup@v4.3.0_6mA@v2 and res_dna_r10.4.1_e8.2_400bps_sup@v4.3.0_4mC_5mC@v1. The latter was obtained from the Rerio repository. MicrobeMod was used to call the methylations both with default settings, and with more stringent criteria i.e., --percent_methylation_cutoff 0.8 and -- methylation_confidence_threshold 0.7.

For PacBio data, the SMRT Analysis pipeline (http://www.pacbiodevnet.com/SMRT-Analysis/Software/SMRT-Pipe) was used for the determination of the epigenetic status of sequenced DNA by identifying the m6A and m4C modified motifs (Clark et al., 2012; Flusberg et al., 2010; Korlach & Turner, 2012) and further curated at REBASE (Roberts et al., 2015).

For the analysis, only motifs detected as methylated by either PacBio or ONT in >80% of their occurrences in the genomes were considered, given that methylation typically occurs in ratios close to 100% in most cases (Hiraoka et al., 2019; Tourancheau et al., 2021).

## Results

### 1. Raw data quality

The details presenting the raw read data quality generated with Illumina, PacBio and ONT across various flow cells and library preparation techniques are given in Tables 1,2, and 3 for TL110, TL19, and TL29, respectively, and in Supplementary Figures S1 and S2.

**Table 1.** Genome assembly results data obtained for the *P. freudenreichii* strain TL110 across different sequencing technologies, library preparations, and assembly strategies.

| Platform | Sequencing strategy (assembler) | contigs | Largest contig | Total length | GC (%) | N50 | Mis assembled contigs | Genome fraction (%)<sup>a</sup> | Mismatch / 100 kbp<sup>b</sup> | Indels / 100 kbp<sup>c</sup> |
| --- | --- | --- | --- | --- | --- | --- | --- | --- | --- | --- |
| Pacbio | (HGAP3) | **1** | **2566312** | **2566312** | **67.33** | **2566312** | **0** | **100.0** | **0.0** | **0.0** |
| Illumina | Nextera (SPAdes) | 127 | 150752 | 2535349 | 67.29 | 43970 | 105 | 98.375 | 0.2 | 0.4 |
| ONT + Illumina | Native R10.4.1 (Unicycler) | 15 | 2227365 | 2565987 | 67.33 | 2227365 | 9 | 99.928 | 4.13 | 8.03 |
|  | Native + BARSEQ R10.4.1 + (Unicycler) | 6 | 2509330 | 2567299 | 67.33 | 2509330 | 5 | 100.0 | 2.92 | 4.83 |
|  | BARSEQ R10.4.1 (Unicycler) | 45 | 544229 | 2501921 | 67.36 | 289201 | 31 | 97.354 | 2.72 | 4.6 |
|  | RAPID R.10.4.1 (Unicycler) | 15 | 2227332 | 2565946 | 67.33 | 2227332 | 10 | 99.928 | 3.82 | 7.05 |
|  | RAPID R9.4.1 (Unicycler) | 4 | 2533496 | 2565690 | 67.32 | 2533496 | 3 | 99.928 | 11.3 | 18.67 |
| ONT | Native R10.4.1 (Flye) | 1 | 2565176 | 2565176 | 67.33 | 2565176 | 0 | 100.0 | 22.03 | 49.0 |
|  | Native + BARSEQ R10.4.1 (Flye) | **1** | **2566302** | **2566302** | **67.33** | **2566302** | **0** | **100.0** | **0.04** | **1.01** |
|  | BARSEQ R10.4.1 (Flye) | 19 | 382593 | 2558590 | 67.33 | 211351 | 18 | 99.327 | 2.38 | 2.03 |
|  | RAPID R.10.4.1 (Flye) | 2 | 2557280 | 2586754 | 67.33 | 2557280 | 2 | 99.946 | 10.09 | 30.97 |

<sup>a,b,c</sup> The comparison has been made against genomes assembled with Pacbio data.

**Table 2.** Genome assembly results data obtained for the *P. freudenreichii* strain TL19 across different sequencing technologies, library preparations, and assembly strategies.

| Platform | Sequencing strategy (assembler) | contigs | Largest contig | Total length | GC (%) | N50 | Mis assembled contigs | Genome fraction (%)<sup>a</sup> | Mismatch / 100 kbp<sup>b</sup> | Indels / 100 kbp<sup>c</sup> |
| --- | --- | --- | --- | --- | --- | --- | --- | --- | --- | --- |
| Pacbio | (HGAP3) | 1 | 2593456 | 2593456 | 67.32 | 2593456 | 0 | 100.0 | 0.0 | 0.0 |
| Illumina | Nextera (Spades) | 158 | 68313 | 2545349 | 67.27 | 31134 | 136 | 97.603 | 0.12 | 3.67 |
| ONT + Illumina | Native R10.4.1 (Unicycler) | 2 | 1660960 | 2497965 | 67.33 | 1660960 | 2 | 96.31 | 1.24 | 5.4 |
|  | Native + BARSEQ R10.4.1 (Unicycler) | 1 | 2497943 | 2497943 | 67.33 | 2497943 | 1 | 96.31 | 1.24 | 5.6 |
|  | BARSEQ R10.4.1 (Unicycler) | 10 | 1419291 | 2470297 | 67.39 | 1419291 | 6 | 95.151 | 0.86 | 5.17 |
|  | RAPID R.10.4.1 (Unicycler) | 1 | 2497944 | 2497944 | 67.33 | 2497944 | 1 | 96.31 | 1.24 | 5.56 |
|  | RAPID R9.4.1 (Unicycler) | 1 | 2497808 | 2497808 | 67.33 | 2497808 | 1 | 96.31 | 1.6 | 9.81 |
| ONT | Native R10.4.1 (Flye) | 1 | 2578969 | 2578969 | 67.33 | 2578969 | 0 | 99.437 | 0.04 | 4.34 |
|  | Native + BARSEQ R10.4.1 (Flye) | 1 | 2578968 | 2578968 | 67.33 | 2578968 | 0 | 99.437 | 0.12 | 4.46 |
|  | BARSEQ R10.4.1 (Flye) | 29 | 303784 | 2584524 | 67.27 | 165525 | 28 | 98.484 | 5.46 | 9.3 |
|  | RAPID R.10.4.1 (Flye) | 1 | 2578974 | 2578974 | 67.33 | 2578974 | 0 | 99.437 | 0.08 | 4.69 |

<sup>a,b,c</sup> The comparison has been made against genomes assembled with Pacbio data.

**Table 3.** Genome assembly results data obtained for the *P. freudenreichii* strain TL29 across different sequencing technologies, library preparations, and assembly strategies.

| Platform | Sequencing strategy (assembler) | contigs | Largest contig | Total length | GC (%) | N50 | Mis assembled contigs | Genome fraction (%)<sup>a</sup> | Mismatch / 100 kbp<sup>b</sup> | Indels / 100 kbp<sup>c</sup> |
| --- | --- | --- | --- | --- | --- | --- | --- | --- | --- | --- |
| Pacbio | (HGAP3) | 2 | 2566309 | 2570072 | 67.3 | 2566309 | 0 | 100.0 | 0.0 | 0.0 |
| Illumina | Nextera (Spades) | 143 | 89206 | 2535202 | 67.28 | 28523 | 125 | 98.167 | 0.04 | 0.51 |
| ONT + Illumina | Native R10.4.1 (Unicycler) | 1 | 2566228 | 2566228 | 67.33 | 2566228 | 0 | 99.854 | 0.58 | 1.83 |
|  | Native + BARSEQ R10.4.1 (Unicycler) | 1 | 2566262 | 2566262 | 67.33 | 2566262 | 0 | 99.854 | 0.35 | 0.66 |
|  | BARSEQ R10.4.1 (Unicycler) | 11 | 1607775 | 2557808 | 67.36 | 1607775 | 7 | 99.419 | 0.31 | 0.63 |
|  | RAPID R.10.4.1 (Unicycler) | 1 | 2566236 | 2566236 | 67.33 | 2566236 | 0 | 99.854 | 0.66 | 1.68 |
|  | RAPID R9.4.1 (Unicycler) | 1 | 2566142 | 2566142 | 67.33 | 2566142 | 0 | 99.854 | 2.73 | 5.26 |
| ONT | Native R10.4.1 (Flye) | 1 | 2565099 | 2565099 | 67.32 | 2565099 | 0 | 99.854 | 20.93 | 52.24 |
|  | Native + BARSEQ R10.4.1 (Flye) | 1 | 2566311 | 2566311 | 67.33 | 2566311 | 0 | 99.854 | 0.19 | 0.78 |
|  | BARSEQ R10.4.1 (Flye) | 22 | 339018 | 2566573 | 67.3 | 174239 | 20 | 99.057 | 1.52 | 1.76 |
|  | RAPID R.10.4.1 (Flye) | 1 | 2565672 | 2565672 | 67.32 | 2565672 | 0 | 99.854 | 9.35 | 30.95 |

<sup>a,b,c</sup> The comparison has been made against genomes assembled with Pacbio data.

No significant differences were observed when comparing the quality for the data obtained utilizing light and dark mode (data not shown), yet an increase of data output was observed.

For V14 chemistry (RAPID and Native and BARSEQ), duplex reads constituted 17-20% of the sequenced bases for the R10.4.1 Native in dark mode (data not shown), as calculated by Dorado. While split pairs did not improve the quality in comparison to regular R10.4.1 data, we observed a drastic increase in the quality score of the paired data compared to regular R10.4.1 Native data (Figure S1f).

### 2. Genome completeness

To assess the performance of diverse library preparation approaches, we examined the genomes of three *Propionibacterium freudenreichii* strains using data from PacBio, Illumina NextSeq 500, and ONT. HGAP3, Spades, and Flye were employed for assembling PacBio, Illumina, and ONT data, respectively. Additionally, a hybrid assembly approach utilizing Unicycler with Illumina short reads was applied to polish ONT assembly data.

The info on *de novo* assembled genomes across different DNA sequencing strategies, DNA library preparation protocols, and assemblers are given in Tables 1-3.

As expected only data from long read sequencing strategies, where DNA library was prepared without PCR and subjected to apt assembler, managed to recover circular single contigs of length > 2.5 Mbp.

Combined data from nanopore BARSEQ and Native barcoding, both sequenced on R10.4.1 and assembled with Flye, gave highest scores in CheckM genome completeness for all three strains (99.5%) compared to all other sequencing strategies. In general, high completeness CheckM scores (> 95%) could be achieved with all sequencing strategies either with Flye, HGAP3, and Spade for ONT, PacBio, and Illumina, respectively, or through hybrid assembly utilizing Unicycler (Figure 3 and supplementary Figure S3).

**Figure 3.**
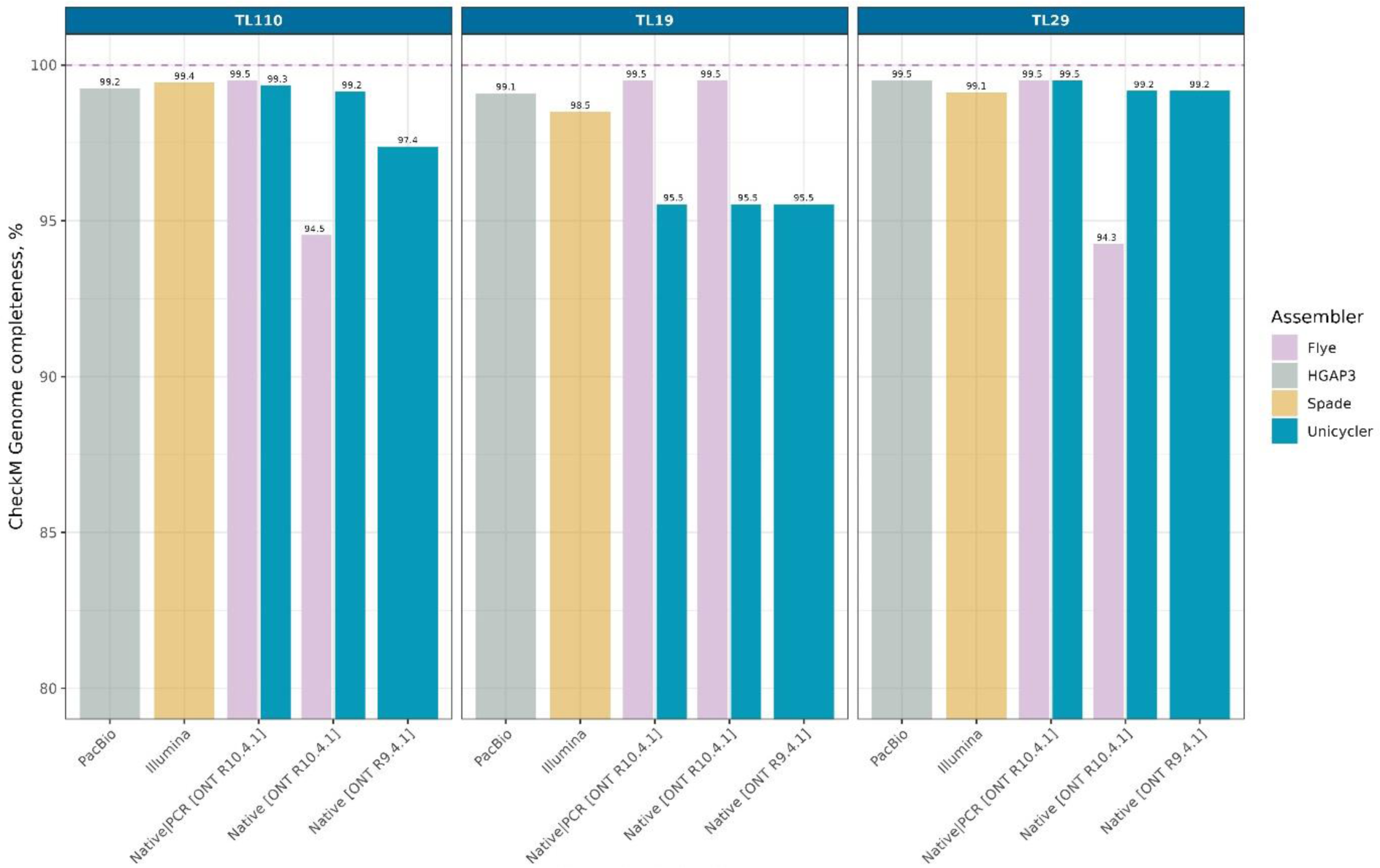
CheckM genome completeness scores for the three *Propionibacterium freudenreichii* strains across different DNA sequencing strategies, library preparation protocols, and assemblers.

The utilization of Illumina short reads for polishing ONT long reads resulted in an increase or decrease in terms of genome completeness, depending on the strain, whereas the utilization of BARSEQ reads to correct long reads always resulted in equal or higher performance.

### 3. Assembled genomes quality: variant calling

PacBio-derived contigs were used as the reference for a variant calling analysis, encompassing insertions, deletions (indels), and substitutions. Among the various variant types examined, deletions had the lowest proportion, followed by insertions, and finally substitutions, across all strains and sequencing strategies tested (Figure 4).

**Figure 4.**
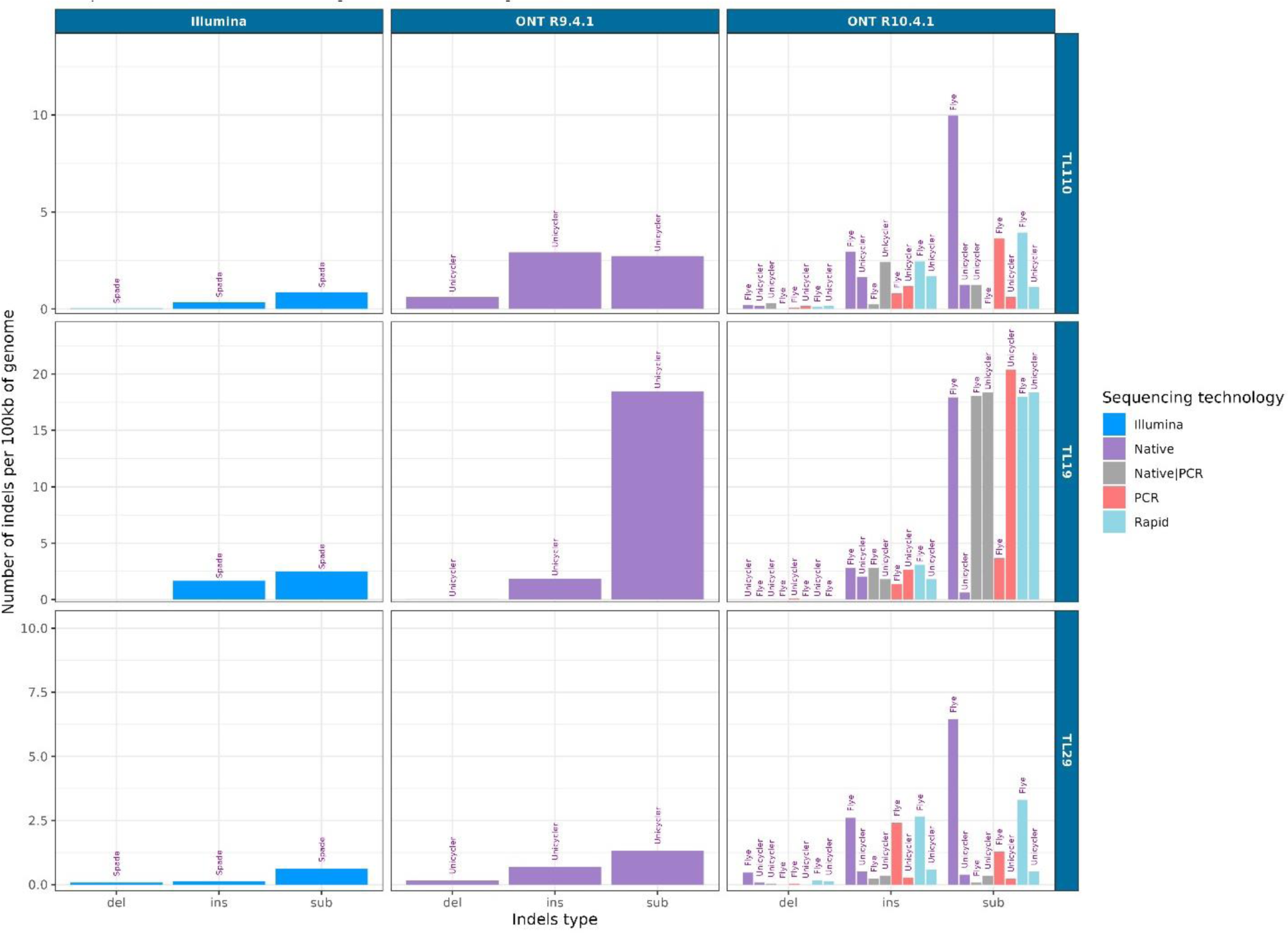
Variant calling for genomes of three strains of *P. freudenreichii*. The comparison was made for assembled genomes recovered using various sequencing technologies and DNA library preparation strategies, and dedicated assemblers. The PacBio data was used as the reference.

The number of deletions was consistently low across three library preparation strategies (BARSEQ, RAPID, and Native) sequenced with the R10.4.1 flow cell. Genomes recovered using data generated with R9.4.1 flow cell, without Illumina read polishing with Unicycler exhibited the highest proportions of insertions, deletions, and substitutions (data not shown). However, hybrid assembly of ONT, R9.4.1 and Illumina data resulted in significant reduction of indels proportion. The lowest proportion of substitutions (lower than that for data generated by Illumina) was obtained with different combinations of library preparations and assembly strategies for data sequenced on ONT R10.4.1.

In general, the best results across three different strains were achieved by combining data from Native barcoding and BARSEQ data generated with R10.4.1 flow cell, assembled with Flye, with one exception where, for the TL19 strain, the number of substitutions remained at 18 substitutions/100Kb, indicating possible strain confounding effect. For DNA libraries prepared with RAPID, BARSEQ, and Native barcoding, hybrid assembly with Unicycler appeared to perform slightly better than Flye, yielding lower proportions of deletions, insertions, and substitutions, except when the data from Native barcoding kit and BARSEQ were combined. No clear differences in the proportions of deletions, insertions, and substitutions were observed between the library preparation strategies (RAPID, Native barcoding, and BARSEQ). The proportions of transitions and transversions across different library preparation strategies and assemblers are provided in supplementary Figure S4.

### 4. Assembled genomes quality: contingency of annotated genes

The assembled genomes of the three *P. freudenreichii* strains were annotated and further evaluated for the proportion of unannotated genes (Figure 4, Figure S3), and prevalence of unique genes (Figure 5) across the different sequencing strategies.

**Figure 5.**
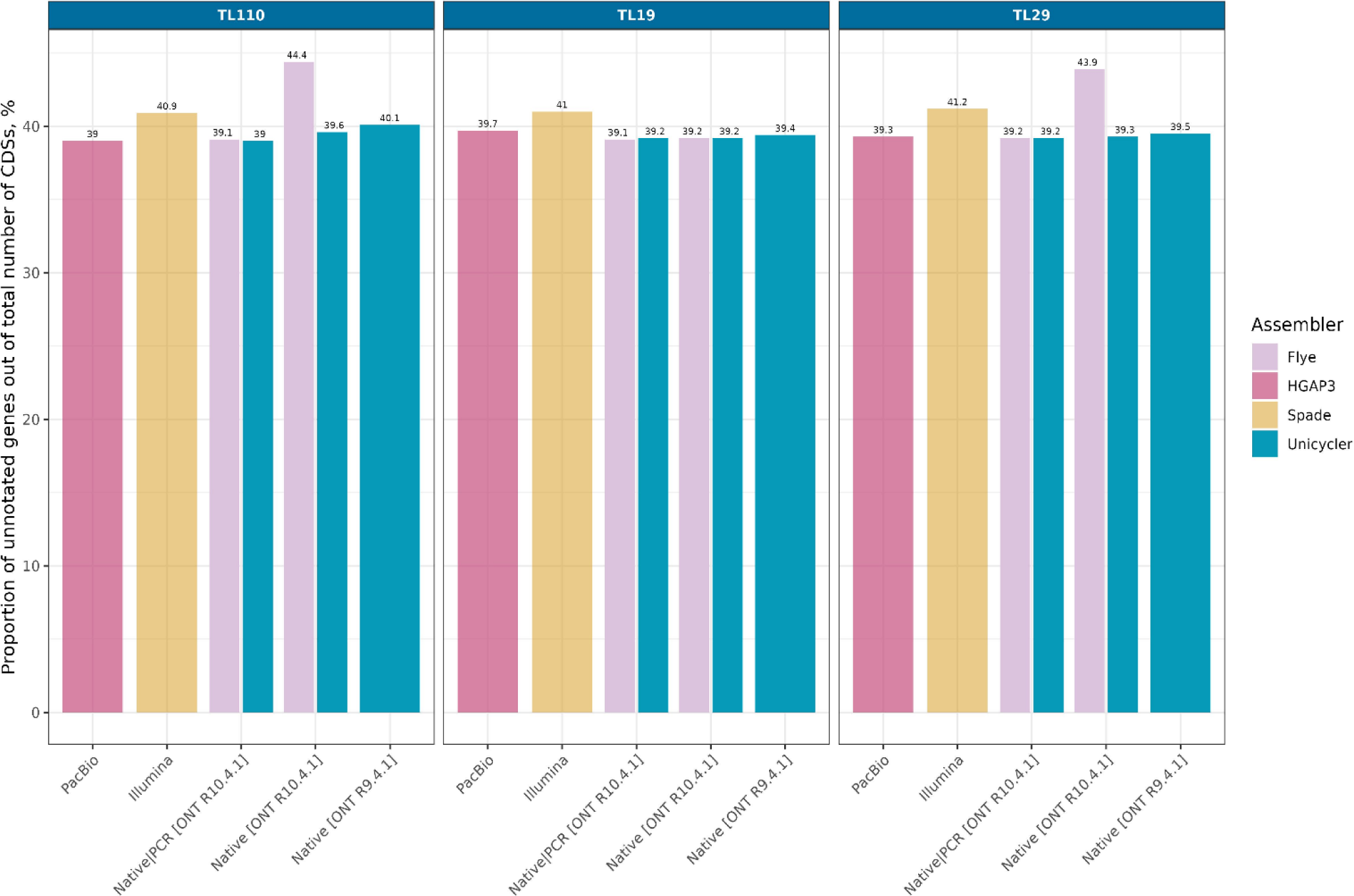
Proportion of unannotated genes in the *P. freudenreichii* strains across different sequencing strategies and assemblers.

Data generated with Illumina and ONT R9.4.1 flow cell resulted in highest proportion of unannotated genes scoring on average 41.0% and 42.5% respectively. The proportion of unannotated genes for all other sequencing strategies ranged between 39% to 40%. The lowest proportions of unannotated genes across the three strains were encountered in data from the Native barcoding kit complemented with BARSEQ data or polished with Illumina data.

Clustering of all genes that demonstrated inconsistency between tested sequencing strategies showed very high level of similarity between the PacBio derived genomes and genomes reconstructed from combined ONT data from Native barcoding kit and BARSEQ sequenced on R10.4.1 flow cell. The results were consistent across all three *P. freudenreichii* strains (Figure 6 and Supplementary Figure S5).

**Figure 6.**
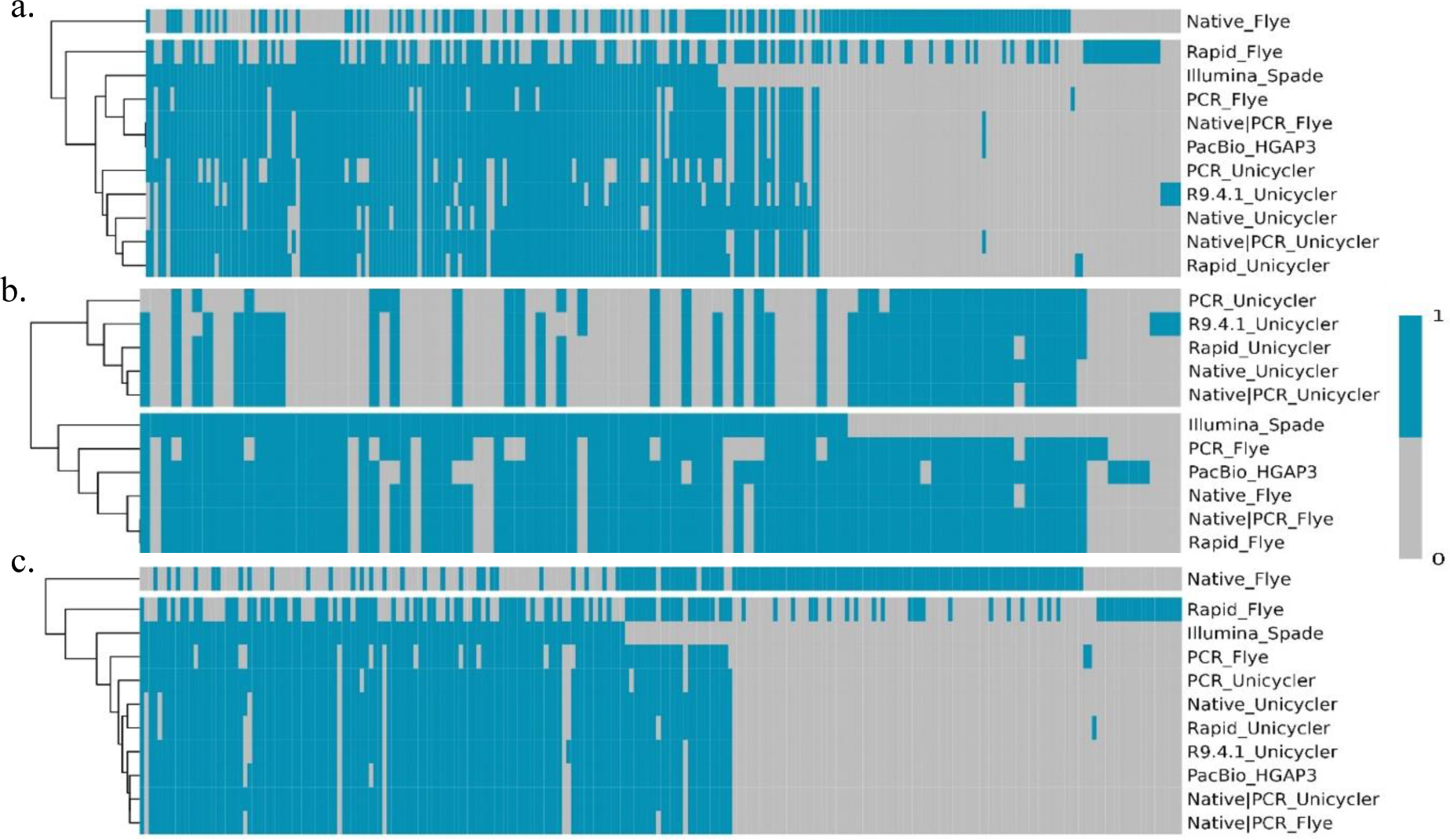
Heatmap demonstrating the contingency of uniquely annotated genes between tested sequencing, library preparation, and assembly strategies for TL110 (a), TL19 (b) and TL29 (c). All genes that were common in all strategies were excluded from the plot.

Interestingly, polishing the ONT R10.4.1 data with Illumina short reads with Unicycler did not consistently lead to achieve results closer to those obtained by PacBio, with some variations observed for the different strains in this regard.

### 5. Comparison of detected methylation motifs across sequencing strategies

The methylation profiles of the three strains were analysed in order to compare the performance of the Native R10.4.1 against the golden standard, PacBio. In general, PacBio outperformed ONT, detecting more reliable motifs i.e., those methylated at or very close to 100 % of the sites with 6mA or 4mC methylations for the three strains (Table 4).

**Table 4.**
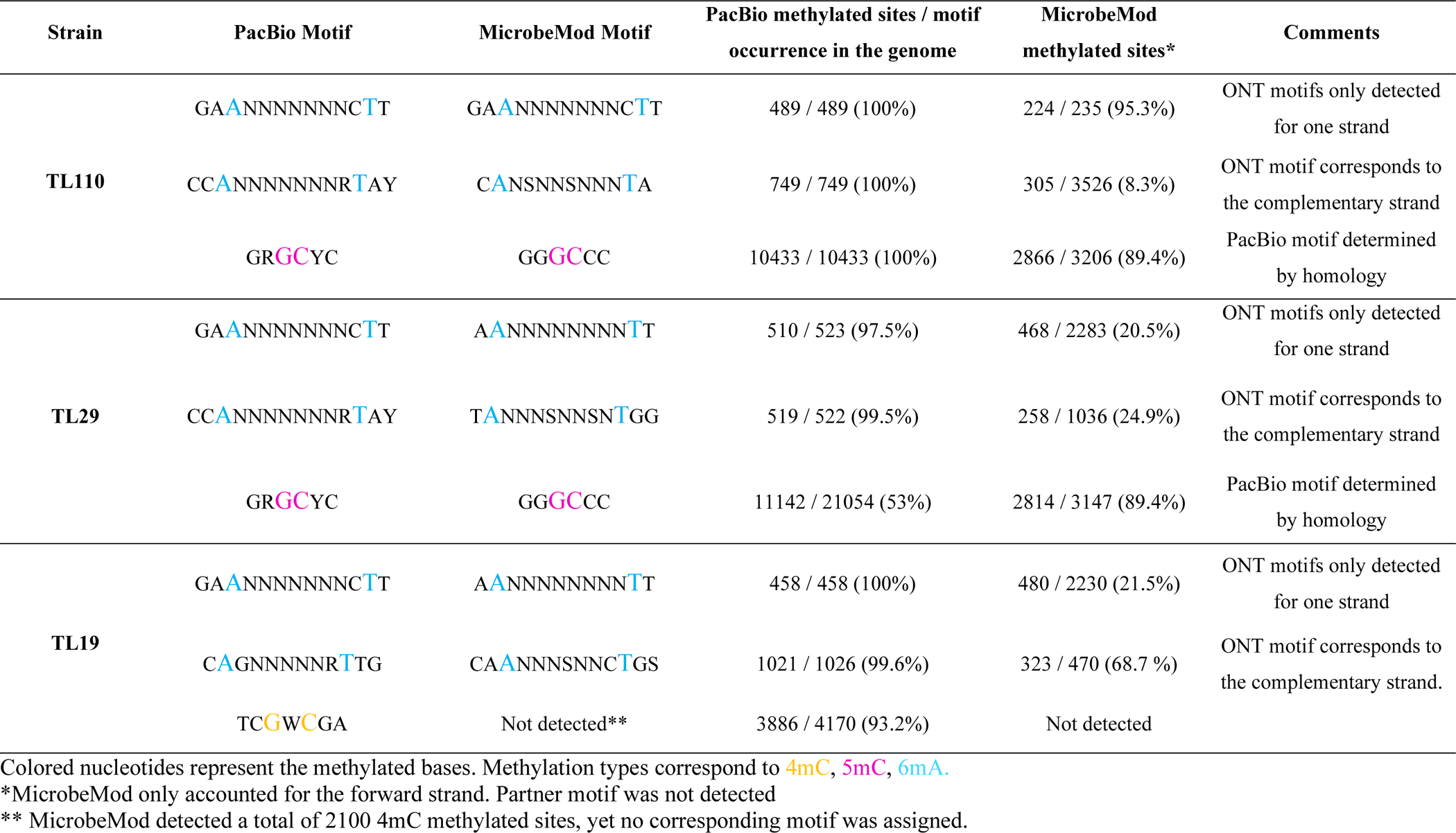
Methylation motifs, types, and occurrence in the genomes of the three *P. freudenreichii* strains obtained with PacBio and ONT R10.4.1 native.

Furthermore, PacBio effectively detected the partner motifs of the type I methyltransferases i.e., the motifs in the complementary strands, a task only partially achieved by MicrobeMod for the GAANNNNNNNCTT motif. The only case in which MicrobeMod performed better than PacBio was for the detection of 5mC methylations for TL110 and TL29 in the motif GGGCCC, with > 2800 methylated sites, representing the 89.4 % of the motifs present in the genome being methylated in both cases, whereas for PacBio the methylation type and motif were assigned based on homology.

## Discussion

Although DNA sequencing has become very cost-effective and high-throughput in recent years, the recovery of complete high-quality genomes remains challenging. It is widely recognized that Illumina platforms generate high-quality data (Hu et al., 2021); however, their short-read nature limits the recovery of complete circular genomes, especially in reconstruction of long repeats across the genome. For this purpose, PacBio is considered the solution as it can produce long reads of good quality that can be assembled into complete genomes (Chin et al., 2013). Oxford Nanopore Technologies (ONT) has been known for many years as the most cost-effective approach with the longest reads, yet historically suffering from a relatively high error rate in assembled genomes (Ip et al., 2015; Laehnemann et al., 2016; Petersen et al., 2019).

Despite recent studies reporting a dramatic improvement in the quality of ONT reconstructed genomes (Sanderson et al., 2023; Sereika et al., 2022), many research groups often opt for other sequencing strategies. Reasons for this choice may include the well-established infrastructure and protocols within a given group, or a lack of clear data on which library preparation strategy should be used to generate high-quality data. For example, Illumina protocols typically require very low concentrations of DNA for library construction, using PCR for enrichment and barcoding. Third-generation sequencing platforms, on the other hand, require higher concentrations, although ONT offers protocols with enrichment using transposase shearing and PCR or MDA.

Sequencing libraries with ONT without enrichment offer several benefits, including the possibility to generate very long reads (Native kits) and the ability to analyze the same data for classical base calling or DNA modification profiles, namely DNA methylation. Although DNA modification calling for bacteria is not yet fully mature on the newest chemistry and flowcells, our data indicate that recovery of methylomes is already feasible using the latest basecaller and methylation calling tools, i.e., dorado v0.5.1 and MicrobeMod (Crits-Christoph et al., 2023.), albeit still lagging behind PacBio. It is clear from our results that the recovery of accurate methylated motifs with ONT R10.4.1 remains challenging, especially when a reference motif is not previously available. It is not unlikely however that newly trained, dedicated baseballers will soon improve recovery of methylation profiles using R10.4.1 flow cell derived data.

Finally, libraries without PCR avoid PCR-related errors, such as primer binding bias, polymerase errors, and transposase-related errors in regions either with high or very low GC content (Benjamini & Speed, 2012; Chen et al., 2013; Sato et al., 2019). Conversely, native DNA often contains modifications which can lead to sequencing errors (Delahaye and Nicolas, 2021), achieving the best base-to-base quality on ONT platforms may require the removal of these modifications with PCR. In our experience, building DNA libraries with the RAPID PCR kit was often unsuccessful and appeared to be highly strain-dependent.

Therefore, we have developed a custom DNA enrichment and barcoding protocol for whole-genome sequencing analysis: Barcode-Amplified Random Sequencing (BARSEQ). In this protocol, DNA is mechanically sheared (using ultrasound or Covaris tubes), followed by ligation of a double-stranded primer binding site on both ends, similar to the Native protocol by ONT. Subsequently, a long-template PCR (3-6 Kb) is performed using primers containing barcodes.

Our data indicate that the newest R10.4.1 flow cells combined with V14 chemistry can recover very high-quality complete bacterial genomes. Our results align with previous studies (Sanderson et al., 2023; Sereika et al., 2022), demonstrating that hybrid assembly using Illumina and ONT data is unnecessary. In fact, our findings even suggest that Unicycler operations on high-quality ONT R10.4.1 data may result in lower quality assemblies when combined with Illumina short-read data.

Although we have not observed clear differences in data quality between RAPID and Native barcoding kits, it is important to mention that Native kits (ligation) typically generate more data than RAPID protocol, which can influence coverage and therefore quality of assembly. BARSEQ consistently produces high-quality data, offering a viable option for scenarios for samples with low DNA concentration. However, it falls short in achieving a satisfactory N50 length for the recovery of complete bacterial genome. It is evident, though, that only by combining long reads from the Native protocol with BARSEQ, data of the highest quality comparable with PacBio can be obtained. Finally, while the duplex reads in the current study contributed to 17-20% of the data with Q25, surpassing the ∼2-6 % reported in other studies (Sanderson et al., 2023), these results are still far from the 30-40% range reported by ONT. Although we observed a considerable increase in read quality when extracting duplex reads and consequently improved assembly compared to Native, assemblies generated with Native protocol combined with BARSEQ remained the most similar to PacBio.

We have estimated the cost (excluding working hours and amortization) of sequencing an average bacterial genome (2.5M) with the described quality using ONT to be approximately $18 (for 96 isolates sequenced on PromethION flow cells), and about 8 to 10 times higher when sequenced with PacBio. This comparison does not include the costs of the equipment that also vary substantially between the two sequencing strategies.

It is essential to note that we have compared all data against genomes recovered with PacBio, as its data has been previously demonstrated to be of the highest quality (Wenger et al., 2019), albeit likely not without any error (Dohm et al., 2020).

It is important to address the main limitations of this study, which include the lack of multiple technical replicates in the experimental design that may limit the generalizability of the findings. Furthermore, the study only tested a limited range of microorganisms, lacking variety in terms of length and GC content, which could affect the applicability of the results to other species. Finally, the absence of plasmids in the tested samples overlooks potential variations in sequencing performance related to extrachromosomal elements, which leaves open windows for further investigations.

## Conclusion

In conclusion, highly comparable high-quality complete bacterial genomes can be reconstructed using ONT with R10.4.1 flow cells and V14 chemistry, comparable to PacBio generated data. The highest quality genomes were reconstructed by combining data from two library preparation strategies: Native barcoding complemented with BARSEQ, both sequenced with ONT. Taking into consideration the significantly lower costs related to the sequencing platform, library preparation, flow cells, and throughput, it appears that ONT is the most optimal sequencing strategy for that task.

## Declaration of interest

All authors believe there is no financial and personal relationships with other people or organizations that could inappropriately influence (bias) their work.

A.F. and R.J.R work for New England Biolabs, a company that sells research reagents, including restriction enzymes and DNA methyltransferases, to the scientific community.

## Acknowledgements

This work was supported by Milk Levy Fund, and co-financed by Arla Foods as a part of the project “Cassandra: Quality modeling using detailed genomics”, and the H2020-MSCA-IF grant nr. 845658.

Data were generated by accessing research infrastructure at University of Copenhagen, including FOODHAY (Food and Health Open Innovation Laboratory, Danish Roadmap for Research Infrastructure).

The authors acknowledge Witold Kot from the University of Copenhagen for the sequencing data of the Illumina samples.

## Data availability

Raw sequences are available at NCBI under the Bioproject accession number PRJNA1070974 and PRJNA772095. SRA accession numbers are included in Supplementary Table 1. All analyses from raw reads to post-assembly quality controls and generated plots were done based on a custom workflow deposited on GitHub (https://github.com/farhadm1990/ONT_helper). The PacBio generated genome sequences are available at NCBI with accession numbers CP085639 for TL29; CP085640 for TL19; CP085641 for TL110.

## References

Achaz, G., Rocha, E. P. C., Netter, P., & Coissac, E. (2002). Origin and fate of repeats in bacteria. In Nucleic Acids Research (Vol. 30, Issue 13).

Bankevich, A., Nurk, S., Antipov, D., Gurevich, A. A., Dvorkin, M., Kulikov, A. S., Lesin, V. M., Nikolenko, S. I., Pham, S., Prjibelski, A. D., Pyshkin, A. V., Sirotkin, A. V., Vyahhi, N., Tesler, G., Alekseyev, M. A., & Pevzner, P. A. (2012). SPAdes: A new genome assembly algorithm and its applications to single-cell sequencing. Journal of Computational Biology, 19(5), 455–477. 10.1089/cmb.2012.0021

Benjamini, Y., & Speed, T. P. (2012). Summarizing and correcting the GC content bias in high-throughput sequencing. Nucleic Acids Research, 40(10). 10.1093/nar/gks001

Boostrom, I., Portal, E. A. R., Spiller, O. B., Walsh, T. R., & Sands, K. (2022). Comparing Long-Read Assemblers to Explore the Potential of a Sustainable Low-Cost, Low-Infrastructure Approach to Sequence Antimicrobial Resistant Bacteria With Oxford Nanopore Sequencing. Frontiers in Microbiology, 13. 10.3389/fmicb.2022.796465

Chen, Y. C., Liu, T., Yu, C. H., Chiang, T. Y., & Hwang, C. C. (2013). Effects of GC Bias in Next-Generation-Sequencing Data on De Novo Genome Assembly. PLoS ONE, 8(4). 10.1371/journal.pone.0062856

Chin, C. S., Alexander, D. H., Marks, P., Klammer, A. A., Drake, J., Heiner, C., Clum, A., Copeland, A., Huddleston, J., Eichler, E. E., Turner, S. W., & Korlach, J. (2013). Nonhybrid, finished microbial genome assemblies from long-read SMRT sequencing data. Nature Methods, 10(6), 563–569. 10.1038/nmeth.2474

Crits-Christoph, A., Kang, S. C., & Lee, H. H. (n.d.). MicrobeMod: A computational toolkit for identifying prokaryotic methylation and restriction-modification with nanopore sequencing. 10.1101/2023.11.13.566931

Dohm, J. C., Peters, P., Stralis-Pavese, N., & Himmelbauer, H. (2020). Benchmarking of long-read correction methods. NAR Genomics and Bioinformatics, 2(2). 10.1093/nargab/lqaa037

Eid, J., Fehr, A., Gray, J., Luong, K., Lyle, J., Otto, G., Peluso, P., Rank, D., Baybayan, P., Bettman, B., Bibillo, A., Bjornson, K., Chaudhuri, B., Christians, F., Cicero, R., Clark, S., Dalal, R., deWinter, A., Dixon, J., … Turner, S. (2009). Real-Time DNA Sequencing from Single Polymerase Molecules. Science, 323(5910), 133–138. 10.1126/science.1162986

Ghazi, A. R., Münch, P. C., Chen, D., Jensen, J., & Huttenhower, C. (2022). Strain Identification and Quantitative Analysis in Microbial Communities. In Journal of Molecular Biology (Vol. 434, Issue 15). Academic Press. 10.1016/j.jmb.2022.167582

Hiraoka, S., Okazaki, Y., Anda, M., Toyoda, A., Nakano, S. ichi, & Iwasaki, W. (2019). Metaepigenomic analysis reveals the unexplored diversity of DNA methylation in an environmental prokaryotic community. Nature Communications, 10(1). 10.1038/s41467-018-08103-y

Hu, T., Chitnis, N., Monos, D., & Dinh, A. (2021). Next-generation sequencing technologies: An overview. Human Immunology, 82(11), 801–811. 10.1016/j.humimm.2021.02.012

Ip, C. L. C., Loose, M., Tyson, J. R., de Cesare, M., Brown, B. L., Jain, M., Leggett, R. M., Eccles, D. A., Zalunin, V., Urban, J. M., Piazza, P., Bowden, R. J., Paten, B., Mwaigwisya, S., Batty, E. M., Simpson, J. T., Snutch, T. P., Birney, E., Buck, D., … Olsen, H. E. (2015). MinION Analysis and Reference Consortium: Phase 1 data release and analysis. F1000Research, 4. 10.12688/f1000research.7201.1

Kolmogorov, M., Yuan, J., Lin, Y., & Pevzner, P. A. (2019). Assembly of long, error-prone reads using repeat graphs. Nature Biotechnology, 37(5), 540–546. 10.1038/s41587-019-0072-8

Laehnemann, D., Borkhardt, A., & McHardy, A. C. (2016). Denoising DNA deep sequencing data-high-throughput sequencing errors and their correction. Briefings in Bioinformatics, 17(1), 154–179. 10.1093/bib/bbv029

Li, H. (2016). Minimap and miniasm: Fast mapping and de novo assembly for noisy long sequences. Bioinformatics, 32(14), 2103–2110. 10.1093/bioinformatics/btw152

Marx, V. (2023). Method of the year: long-read sequencing. In Nature Methods (Vol. 20, Issue 1, pp. 6–11). Nature Research. 10.1038/s41592-022-01730-w

Parks, D. H., Imelfort, M., Skennerton, C. T., Hugenholtz, P., & Tyson, G. W. (2015). CheckM: Assessing the quality of microbial genomes recovered from isolates, single cells, and metagenomes. Genome Research, 25(7), 1043–1055. 10.1101/gr.186072.114

Petersen, L. M., Martin, I. W., Moschetti, W. E., Kershaw, C. M., & Tsongalis, G. J. (2019). Third-Generation Sequencing in the Clinical Laboratory: Exploring the Advantages and Challenges of Nanopore Sequencing. https://journals.asm.org/journal/jcm

Sanderson, N. D., Kapel, N., Rodger, G., Webster, H., Lipworth, S., Street, T. L., Peto, T., Crook, D., & Stoesser, N. (2023). Comparison of R9.4.1/Kit10 and R10/Kit12 Oxford Nanopore flowcells and chemistries in bacterial genome reconstruction. Microbial Genomics, 9(1). 10.1099/mgen.0.000910

Sato, M. P., Ogura, Y., Nakamura, K., Nishida, R., Gotoh, Y., Hayashi, M., Hisatsune, J., Sugai, M., Takehiko, I., & Hayashi, T. (2019). Comparison of the sequencing bias of currently available library preparation kits for Illumina sequencing of bacterial genomes and metagenomes. DNA Research, 26(5), 391–398. 10.1093/dnares/dsz017

Seemann, T. (2014). Prokka: Rapid prokaryotic genome annotation. Bioinformatics, 30(14), 2068–2069. 10.1093/bioinformatics/btu153

Sereika, M., Kirkegaard, R. H., Karst, S. M., Michaelsen, T. Y., Sørensen, E. A., Wollenberg, R. D., & Albertsen, M. (2022). Oxford Nanopore R10.4 long-read sequencing enables the generation of near-finished bacterial genomes from pure cultures and metagenomes without short-read or reference polishing. Nature Methods, 19(7), 823–826. 10.1038/s41592-022-01539-7

Siguier, P., Perochon, J., Lestrade, L., Mahillon, J., & Chandler, M. (2006). ISfinder: the reference centre for bacterial insertion sequences. Nucleic Acids Research, 34(Database issue). 10.1093/nar/gkj014

Tourancheau, A., Mead, E. A., Zhang, X. S., & Fang, G. (2021). Discovering multiple types of DNA methylation from bacteria and microbiome using nanopore sequencing. Nature Methods, 18(5), 491–498. 10.1038/s41592-021-01109-3

Vaser, R., Sović, I., Nagarajan, N., & Šikić, M. (2017). Fast and accurate de novo genome assembly from long uncorrected reads. Genome Research, 27(5), 737–746. 10.1101/gr.214270.116

Vilella (2023) New Oxford Nanopore ‘dark mode’ sequencing. Albert Vilella’s Rhymes with Haystack. https://albertvilella.substack.com/p/new-oxford-nanopore-dark-mode-sequencing

Wang, Y., Zhao, Y., Bollas, A., Wang, Y., & Au, K. F. (2021). Nanopore sequencing technology, bioinformatics and applications. In Nature Biotechnology (Vol. 39, Issue 11, pp. 1348–1365). Nature Research. 10.1038/s41587-021-01108-x

Wick, R. R., Judd, L. M., Gorrie, C. L., & Holt, K. E. (2017). Unicycler: Resolving bacterial genome assemblies from short and long sequencing reads. PLoS Computational Biology, 13(6). 10.1371/journal.pcbi.1005595

Zhao, W., Zeng, W., Pang, B., Luo, M., Peng, Y., Xu, J., Kan, B., Li, Z., & Lu, X. (2023). Oxford nanopore long-read sequencing enables the generation of complete bacterial and plasmid genomes without short-read sequencing. Frontiers in Microbiology, 14. 10.3389/fmicb.2023.1179966

