## Supplementary Figures and Table for "Matching Excellence: ONT’s Rise to Parity with PacBio in Genome Reconstruction of Non-Model Bacterium with High GC Content"

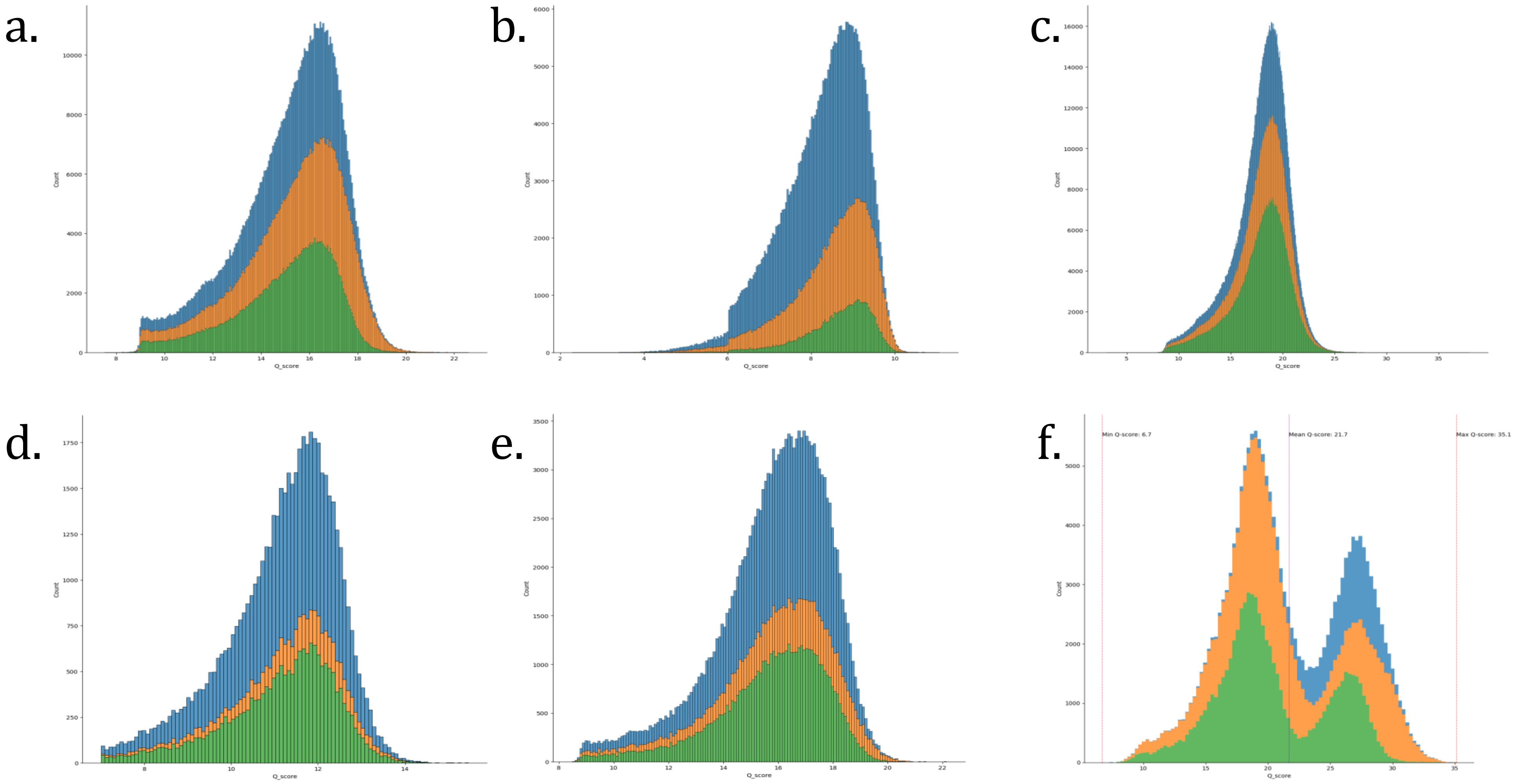


**Figure S1.** Quality histograms of the raw data for PacBio and ONT reads, in which (a) ONT R10.4.1 native; (b) PacBio; (c) R10.4.1 shotgun PCR; (d) ONT R9.4.1 native; (e) ONT R10.4.1 RAPID.

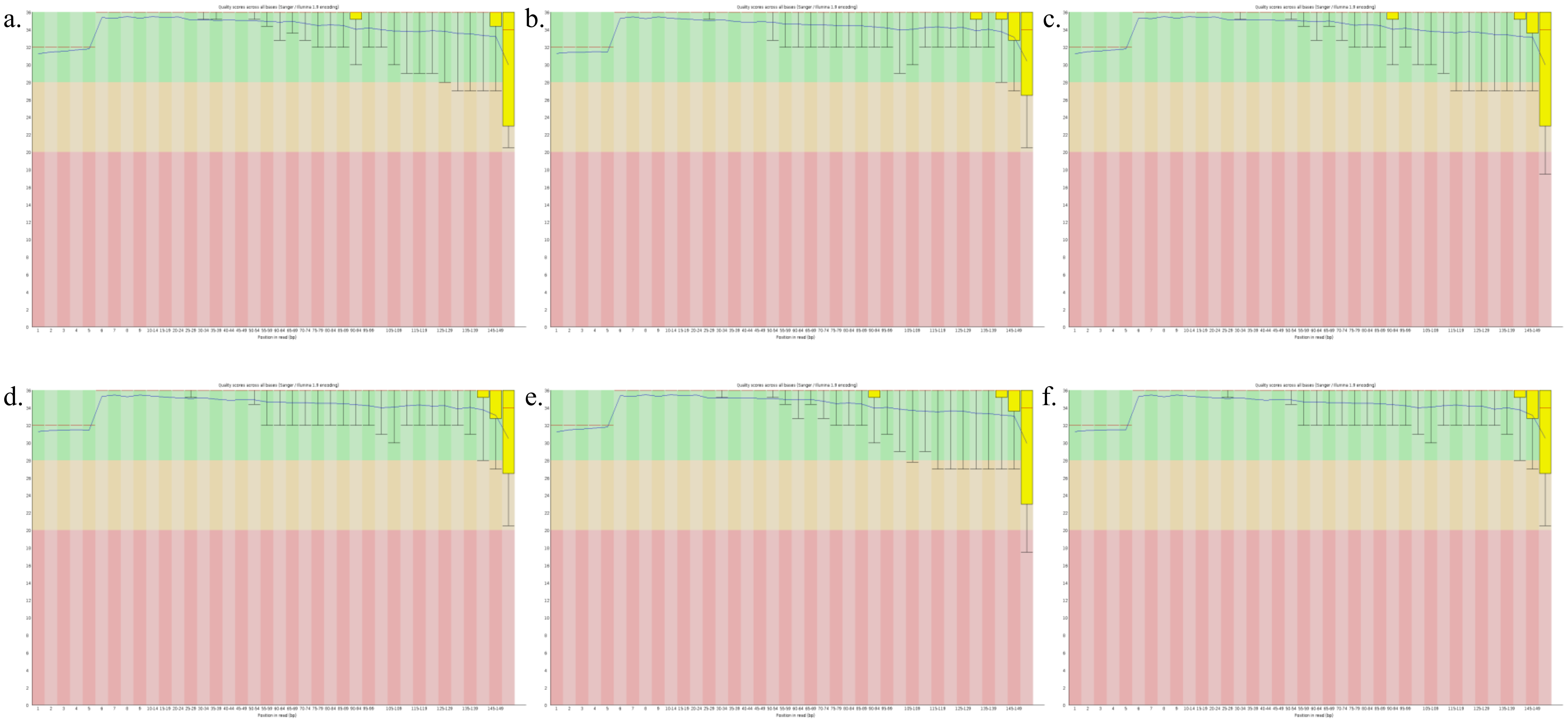

**Figure S2.** Quality scores for the Illumina reads, in which (a) TL110 R1; (b) TL110 R2; (c) TL29 R1; (d) TL29 R2; (e) TL19 R1; (f) TL19 R2.

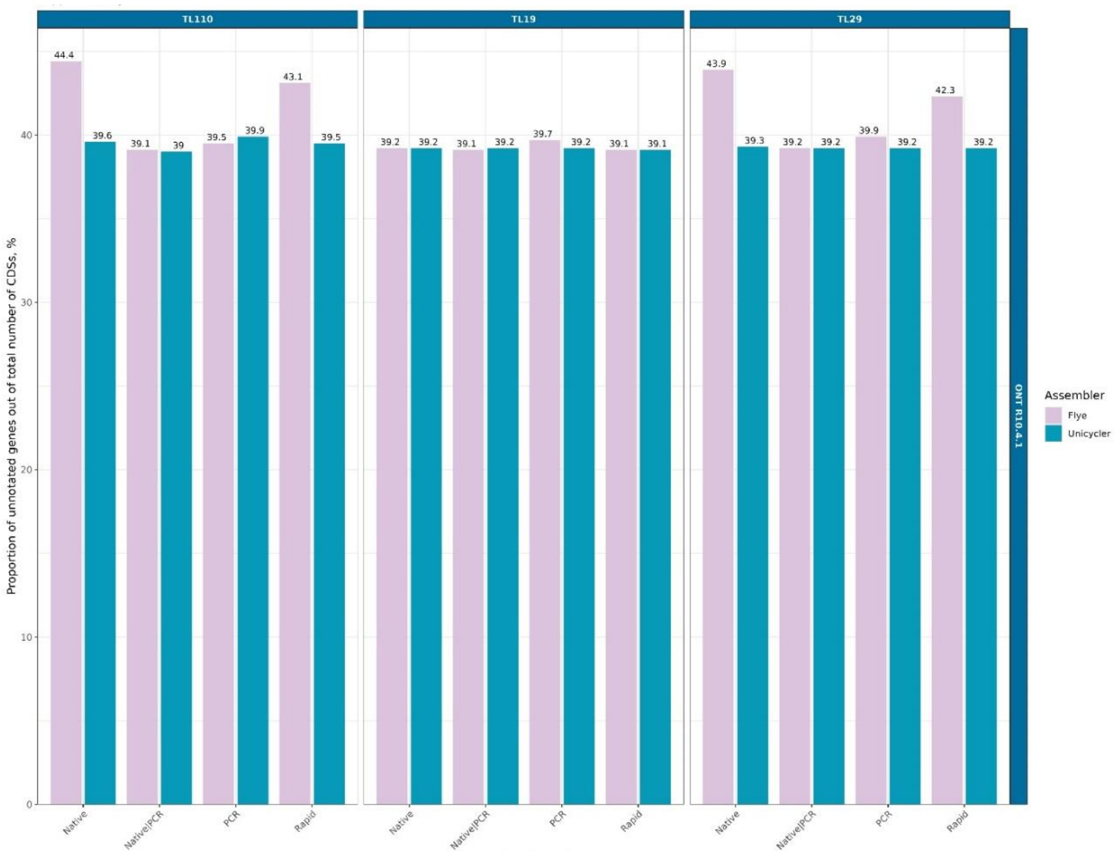


**Figure S3.** Proportion of unannotated genes in ONT R10.4.1 with the different library preparation strategies.

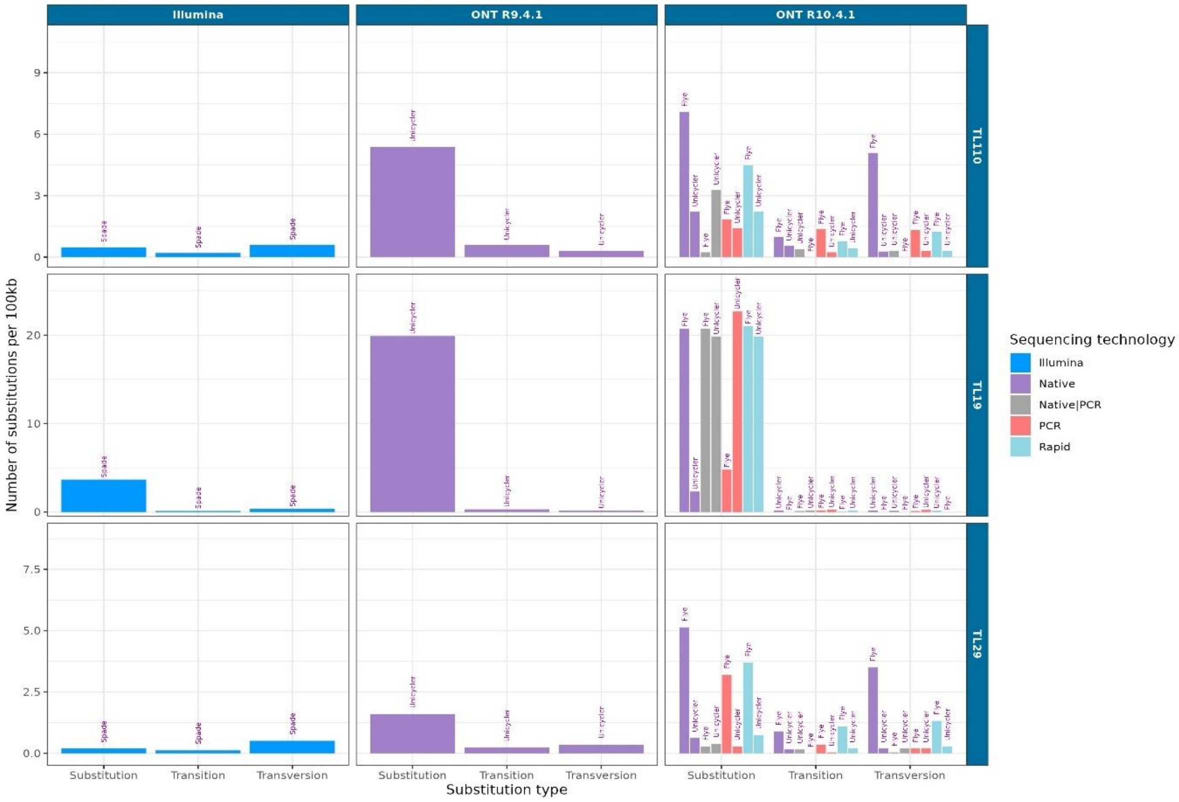


**Figure S4.** Substitution types per 100kbp for the different sequencing strategies. Comparison was made based on alignment to the PacBio reference genome.

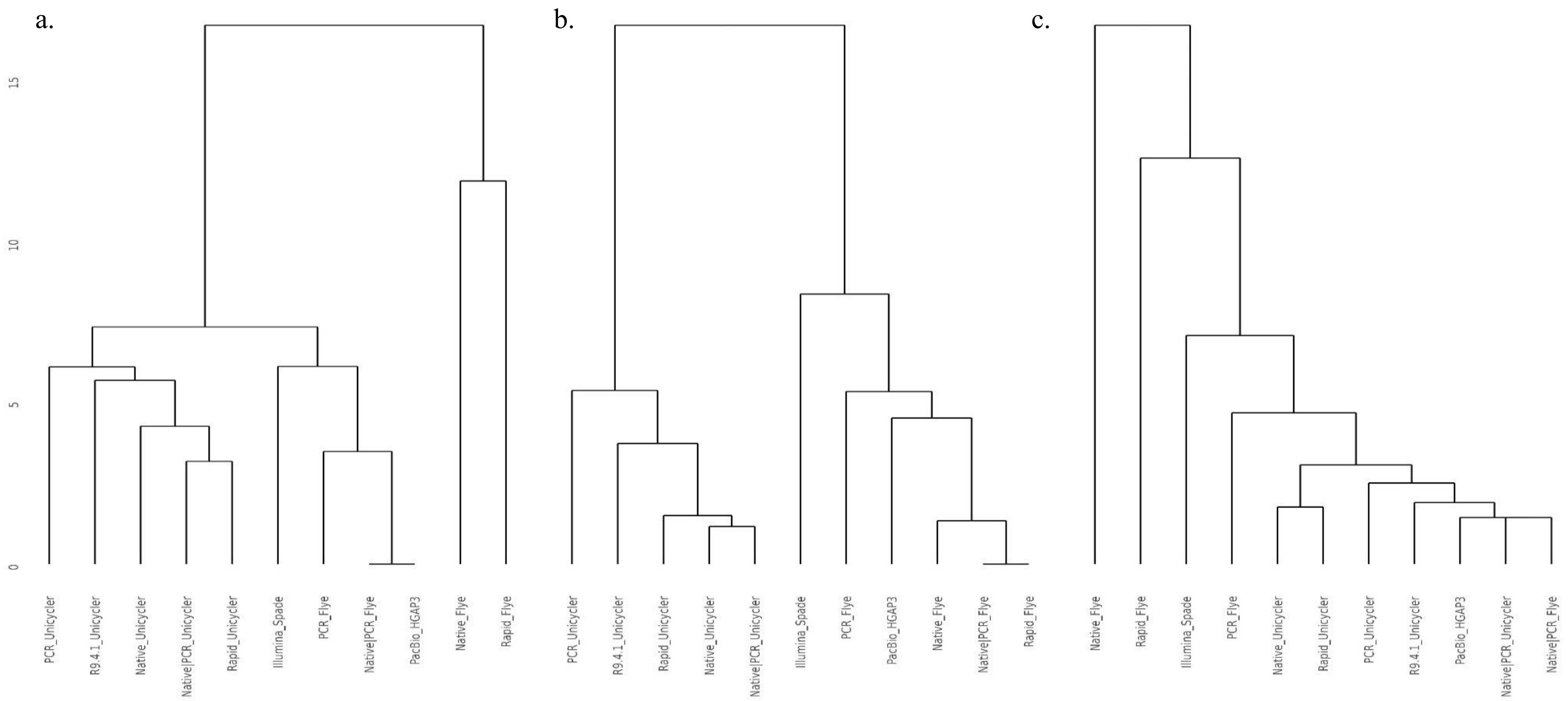


**Figure S5.** Dendrogram of ward.D2 distance regarding annotated genes for all sequencing strategies, in which a) TL110; b) TL19; c) TL29. For the analysis, all the shared genes were excluded.

**Supplementary Table 1.** Accession numbers for strains TL110, TL19, and TL29 across different sequencing strategies.

| Isolate | Library preparation | Flowcell | SRA accession no. |
|---|---|---|---|
| TL110 | ONT Native (dark) | R10.4.1 | SRR27779107 |
| TL110 | ONT Native (light) | R10.4.1 | SRR27779106 |
| TL110 | ONT Rapid | R9.4.1 | SRR27779095 |
| TL110 | ONT Rapid | R10.4.1 | SRR27779097 |
| TL110 | ONT BARSEQ | R10.4.1 | SRR27779096 |
| TL110 | Illumina | - | SRR27779094 |
| TL110 | PacBio | - | SRS10712865 |
| TL19 | ONT Native (dark) | R10.4.1 | SRR27779093 |
| TL19 | ONT Native (light) | R10.4.1 | SRR27779092 |
| TL19 | ONT Rapid | R9.4.1 | SRR27779105 |
| TL19 | ONT Rapid | R10.4.1 | SRR27779091 |
| TL19 | ONT BARSEQ | R10.4.1 | SRR27779090 |
| TL19 | Illumina | - | SRR27779104 |
| TL19 | PacBio | - | SRS10717466 |
| TL29 | ONT Native (dark) | R10.4.1 | SRR27779103 |
| TL29 | ONT Native (light) | R10.4.1 | SRR27779102 |
| TL29 | ONT Rapid | R9.4.1 | SRR27779099 |
| TL29 | ONT Rapid | R10.4.1 | SRR27779101 |
| TL29 | ONT BARSEQ | R10.4.1 | SRR27779100 |
| TL29 | Illumina | - | SRR27779098 |
| TL29 | PacBio | - | SRS10725002 |
